# Postnatal Development Of Dendritic Morphology And Action Potential Shape In Rat Substantia Nigra Dopaminergic Neurons

**DOI:** 10.1101/2024.09.18.613620

**Authors:** Estelle Moubarak, Florian Wernert, Fabien Tell, Jean-Marc Goaillard

## Abstract

Substantia nigra pars compacta (SNc) dopaminergic (DA) neurons are characterized by specific morphological and electrophysiological properties. First, in ∼90% of the cases, their axon arises from an axon-bearing dendrite (ABD) at highly variable distances from the soma. Second, they display a highly regular pattern of spontaneous activity (aka pacemaking) and a broad action potential (AP) that faithfully back-propagate through the entire dendritic arbor. In previous studies (Moubarak et al., 2019; Moubarak et al., 2022), we demonstrated that the presence of a high density of sodium current in the ABD and the complexity of this dendrite played a critical role in the robustness of pacemaking and setting the half-width of the AP. In the current study, we investigated the postnatal development of both morphology and AP shape in SNc DA neurons in order to determine when and how the mature electrophysiological phenotype of these neurons was achieved. To do so, we performed electrophysiological recordings of SNc DA neurons at 4 postnatal ages (P3, P7, P14, P21) and fully reconstructed their dendritic and proximal axon morphology. Our results show that several morphological parameters, including the length of the ABD, display abrupt changes between P7 and P14, such that a mature morphology is reached by P14. We then showed that AP shape followed a similar timecourse. Using realistic multicompartment Hodgkin-Huxley modeling, we then demonstrated that the rapid morpho-electrical maturation of SNc DA neurons likely arises from synergistic increases in dendritic length and in somatodendritic sodium channel density.

**Significance statement:** Substantia nigra pars compacta (SNc) dopaminergic (DA) neurons display several morphological and electrophysiological peculiarities. For instance, their axon arises in most cases from an axon-bearing dendrite (ABD) and their action potential (AP) is broad and faithfully back-propagates through the entire dendritic tree. In the present study, we performed electrophysiological recordings, neuronal reconstruction and computational modeling to determine the postnatal development of dendritic morphology and AP shape in SNc DA neurons. We found that ABD length rapidly increases after post-natal day 7 (P7) to reach maturity by P14 and that AP shape follows a similar timecourse. Computational modeling then suggested that the achievement of a mature AP comes from synergistic increases in dendritic length and in somatodendritic sodium channel density.

## Introduction

One of the defining features of neurons compared to other cell types is their large size and complex morphology, each neuron harboring one long axon and usually several primary dendrites arising from the cell body. Across neuronal types, morphology greatly varies, such that it constitutes one of the first criteria used to identify and distinguish neuronal types (Migliore and Shepherd, 2005; Ascoli et al., 2008; Gouwens et al., 2020): in particular, the number of dendrites, their orientation, branching pattern and the presence/absence of dendritic spines are defining factors of each neuronal type (y Cajal and Azoulay, 1952). Variations in dendritic morphology have a strong influence on neuronal function and are associated with differences in the propagation of electrical signals (Rall, 1969; Rinzel and Rall, 1974; Vetter et al., 2001). For instance, very complex dendritic trees such as those found in cerebellum Purkinje cells promote a passive propagation of signals while the much simpler dendrites of substantia nigra pars compacta (SNc) dopaminergic (DA) neurons facilitate the faithful back-propagation of action potentials (APs) (Vetter et al., 2001).

This latter neuronal type has another distinctive morphological feature: in ∼90% of mature SNc DA neurons, the axon arises from a dendrite (hence called the axon-bearing dendrite, or ABD) and not from the soma (Grace and Bunney, 1983; Hausser et al., 1995; Moubarak et al., 2019). In fact, it was demonstrated that the axon is located at highly variable distances from the soma that can exceed 200µm in adult rat neurons (Hausser et al., 1995). In two previous studies, we investigated whether this morphological peculiarity could influence the activity of SNc DA neurons, in particular their ability to generate an autonomous regular tonic pattern of activity (aka pacemaking) and faithfully back-propagate APs (Moubarak et al., 2019; Moubarak et al., 2022). Extending previous work from other groups (Wilson and Callaway, 2000; Jang et al., 2014), we demonstrated that the presence of a high density of sodium channels in the somato-dendritic compartment was not only responsible for the faithful back-propagation of the APs but also sustained pacemaking and allowed it to be insensitive to large variations in axon location (Moubarak et al., 2019). We also showed that the ABD of mature SNc DA neurons was on average much longer and more complex (composed of more segments) than the non-axon bearing dendrites (nABDs), although this particular dendritic topology displayed a high degree of cell-to-cell variability (Moubarak et al., 2022). In addition, we could show that these variations in dendritic topology were associated with variations in half-width of the somatically recorded AP, such that the ABD seemed to accelerate the AP while nABDs slowed it down. This last study suggested that the opposite influences of the ABD and nABDs were supported by a higher density of sodium channels in the ABD. Both studies highlighted the critical role played by the ABD in the mature electrophysiological phenotype of SNc DA neurons, essentially due to the expression of a high density of sodium channels in this dendritic compartment (Moubarak et al., 2019; Moubarak et al., 2022).

Interestingly, in another study, we demonstrated that the mature electrophysiological phenotype of rat SNc DA neurons (in particular their regular pacemaking behavior) is acquired over the first three post-natal weeks, with two main transitions along this developmental time course (Dufour et al., 2014): while SNc DA neurons display a bursting pattern of activity at very early stages (P2-P3), their activity evolves towards an irregular pattern (between P5 and P10) before becoming tonic and regular (after P14). This change in firing pattern is associated with modifications in the amplitude of the AP and the afterhyperpolarization (AHP), suggesting that the densities of sodium channels and calcium-activated potassium channels (SK) might steeply increase after P5 to reach their mature state at P14 (Dufour et al., 2014). Hence, although P14 SNc DA neurons cannot be considered “adult”, their electrophysiological phenotype barely changes after this developmental stage.

Because of ***i)*** the demonstrated implication of somato-dendritic sodium channels in pacemaking and action potential ***ii)*** the particular role of the ABD in sustaining and shaping the back-propagating AP and ***iii)*** the particular dendritic topology observed in mature SNc DA neurons (ABD longer and more complex than nABDs), we wondered whether the peculiar developmental trajectory of the electrophysiological phenotype of SNc DA neurons (Dufour et al., 2014) could be associated with changes in dendritic morphology. To address this question, we performed patch-clamp recordings of neurobiotin-filled SNc DA neurons from P3, P7, P14 and P21 rats and fully reconstructed their morphology. In addition, we used realistic multi-compartment Hodgkin-Huxley modeling to investigate the respective roles of morphology and biophysical properties in the development of the electrophysiological phenotype of this neuronal type.

## Material and Methods

The dataset of P21 neurons (n=58) used in the current study is partly the same that was used in two previous publications (Moubarak et al., 2019; Moubarak et al., 2022) and was used here as a comparison with the new datasets obtained from P3 (n=14 neurons), P7 (n=35) and P14 rats (n=16) in order to provide a description of postnatal morphological development.

### Acute midbrain slice preparation

Acute slices were prepared from P2-P3 (mean=2.4, N=5), P6-P8 (mean=6.9, N=10), P13-P16 (mean=14.7, N=7) and P19-P21 (mean=P20, N=12) Wistar rats of either sex. All experiments were performed according to the European (Council Directive 86/609/EEC) and institutional guidelines for the care and use of laboratory animals (French National Research Council). Rats were anesthetized with isoflurane (CSP) in an oxygenated chamber (TEM SEGA) and decapitated. The brain was immersed briefly in oxygenated ice-cold low-calcium artificial cerebrospinal fluid (aCSF) containing the following (in mM): 125 NaCl, 25 NaHCO_3_, 2.5 KCl, 1.25 NaH_2_PO_4_, 0.5 CaCl_2_, 4 MgCl_2_, 25 D-glucose, pH 7.4, oxygenated with 95% O_2_/5% CO_2_ gas. The cortices were removed and coronal midbrain slices (250μm) were cut on a vibratome (Leica VT 1200S) in oxygenated ice-cold low calcium aCSF. Following 20-30 min incubation in 32°C oxygenated low calcium aCSF, the slices were incubated for at least 30 minutes in oxygenated aCSF (125NaCl, 25 NaHCO_3_, 2.5 KCl, 1.25 NaH_2_PO_4_, 2 CaCl_2_, 2 MgCl_2_ and 25 glucose, pH 7.4, oxygenated with 95% O_2_ 5% CO_2_ gas) at room temperature prior to electrophysiological recordings.

### Drugs

Picrotoxin (100µM, Sigma Aldrich) and kynurenate (2mM, Sigma Aldrich) were bath-applied via continuous perfusion in aCSF to block inhibitory and excitatory synaptic activity, respectively.

### Electrophysiology recordings and analysis

All recordings (n=123 cells from N=34 rats) were performed on midbrain slices continuously superfused with oxygenated aCSF. Patch pipettes (1.8-4 MOhm) were pulled from borosilicate glass (GC150TF-10, Havard Apparatus) on a DMZ Universal Puller (Zeitz Instruments). The patch solution contained in mM: 20KCl, 10 HEPES, 10 EGTA, 2 MgCl2, 2 Na-ATP and 120 K-gluconate, pH 7.4, 290-300 mOsm. Neurobiotin (0.05%; Vector Labs) was included in the intracellular solution to allow morphological reconstruction and identification of dopaminergic neurons using post-hoc tyrosine-hydroxylase immunolabeling (Moubarak et al., 2019). Whole-cell recordings were made from SNc dopaminergic neurons visualized using infrared differential interference contrast videomicroscopy (QImaging Retiga camera; Olympus BX51WI microscope) and identified as previously described (Moubarak et al., 2019). Whole-cell current-clamp recordings with a series resistance <10MOhm (soma) were included in the study. Capacitive currents and liquid junction potential (+13.2mV) were compensated online and offset potentials were measured after removing the pipette from the neuron. Bridge balance (100%, 10µs) was used to compensate series resistance. Recordings with offset values above 1mV were discarded from the analysis. Recordings were acquired at 50kHz and were filtered with a low-pass filter (Bessel characteristic between 2.9 and 5kHz cutoff frequency). Action potentials (APs) generated during a 40s period of spontaneous activity were averaged and the amplitude and duration of the AP at half of the maximal height of the AP (AP half-width) were measured. AP threshold was measured as the voltage value corresponding to the crossing of a 20mV/ms voltage slope on the first time-derivative of voltage. The second time-derivative of voltage was used to measure the amplitudes of the initial segment (IS) and somatodendritic (SD) components of the AP.

### Electrophysiology data acquisition and analysis

Data were acquired with a HEKA EPC 10/USB patch-clamp amplifier (HEKA electronics) and patchmaster software (HEKA electronics) or a Multiclamp 700B (Molecular Devices, Palo Alto, CA). Analysis was conducted using FitMaster v2×30 (Heka Elektronik) and Clampfit (Molecular Devices).

### Immunohistochemistry and morphological reconstruction

Acute slices containing Neurobiotin-filled cells were fixed 30 min in 4% paraformaldehyde at 4°C and immunolabelled with anti-tyrosine hydroxylase (chicken polyclonal, Abcam, 1:1000) and anti-ankyrinG (mouse monoclonal IgG2b, NeuroMab, 1:250) antibodies. Goat anti-mouse IgG2b Alexa Fluor 488 (Invitrogen; 1:1000; 2µg/ml) and goat anti-chicken Alexa Fluor 633 (Invitrogen; 1:3000; 1.66ng/ml) were used to reveal tyrosine hydroxylase and ankyrinG stainings, respectively. Streptavidin Alexa Fluor 594 (Invitrogen; 1:12000; 1.66ng/ml) was used to reveal neurobiotin labelling. Slices were mounted in Faramount mounting medium (Dako). Immunolabellings were viewed on an LSM 780 Zeiss (Carl Zeiss AG, Germany) and images were captured using Zeiss ZEN software. Images were analyzed with Fiji/ImageJ software (Schindelin et al., 2012; Rueden et al., 2017) and neurons were reconstructed using the Simple Neurite Tracer plugin (Longair et al., 2011). The axon was identified using ankyrinG labelling, which allowed us to discriminate between ABD and nABDs. All dendritic lengths were extracted directly from the paths traced through the stack images of the neurons. To define dendritic complexity, we extracted from the reconstructions the number of dendritic segments (branches) found on the ABD and nABDs. The average length and number of segments per nABD were obtained by dividing the total for these parameters by the number of primary nABD dendrites for each neuron. Soma volumes were estimated by using the “fill out path” method on Simple Neurite Tracer.

### Multi-compartment modelling

Simulations were performed using NEURON 7.5 software (Hines and Carnevale, 2001), as described in two previous studies (Moubarak et al., 2019; Moubarak et al., 2022). Neuronal mophologies corresponding to the realistic morphologies from 19 reconstructed P7 and 36 P21 SNc DA neurons were used. The 36 P21 models correspond to data already published (Moubarak et al., 2019; Moubarak et al., 2022) that were used for comparison with the P7 data obtained for the present study.

For each compartment, membrane voltage was obtained as the time integral of a first-order differential equation:

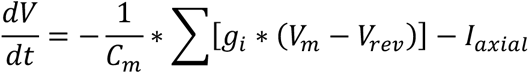

where *V_m_* is the membrane potential, *C_m_* the membrane capacitance, *g_i_* are ionic conductances and *V_rev_* their respective reversal potentials. The axial flow of current (*I_axial_*) between adjacent compartments is calculated by the NEURON simulation package (Hines and Carnevale, 2001). Cytoplasmic resistivity, specific membrane capacitance and specific membrane resistance were set to 150 Ohm.cm, 0.75 µF/cm², and 100,000 Ohm*cm², respectively, with the reversal potential for the leak conductance set at -50 mV. Active conductances followed activation-inactivation Hodgkin-Huxley kinetics (see (Moubarak et al., 2019; Moubarak et al., 2022)).

Parameters for I_A_, I_CaL_, I_KCa_ and I_H_ were based on previous published values for SNc DA neurons (Wilson and Callaway, 2000; Amendola et al., 2012; Engel and Seutin, 2015). Fast sodium and potassium currents were derived from Migliore and Schild models, respectively (Schild et al., 1993; Migliore et al., 2008). The SK current is solely activated by an increase in calcium concentration. Therefore, intracellular calcium uptake was modeled as a simple decaying model according to (Destexhe et al., 1993). Conductance values were set according to our own measurements or published values (see (Moubarak et al., 2019; Moubarak et al., 2022)). Consistent with the literature, g_Na_ and g_KDR_ densities are higher in the AIS than in the rest of the neuron so that AP always initiates in the AIS (Zhou et al., 1998; Kole et al., 2008; Hu et al., 2009; Gonzalez-Cabrera et al., 2017). According to Gentet and Williams, I_A_ density and inactivation kinetics were higher and depolarized, respectively, in the soma compared to the dendritic arbor (Gentet and Williams, 2007). Initializing potential was set at -70 mV and analysis was performed after pacemaking frequency reached a steady-state (8 spikes). Each simulation run had a 6000 ms duration with a dt of 0.01 ms. All dendritic compartments and the axon-start compartment contained all currents whereas AIS and axon only contained fast sodium and potassium currents (g_Na_, g_KDR_). Unless otherwise stated, all currents but the fast sodium and calcium currents had fixed and homogeneous conductance values in the dendrites and the axon-start compartment.

For the realistic morphology models, exact dendrite lengths, soma volume, and diameters of primary dendrites, ABD secondary dendrites, axon and AIS were used (see the ***Immunohistochemistry and morphological reconstruction*** section for details). The specific branching patterns of each neuron (topology) were also respected. In order to be consistent with the NEURON software constraints, soma volume was implemented by computing the equivalent cylinder corresponding to the volume measured using “fill out path” method in Simple Neurite Tracer. Axonal diameter was considered constant and set to 0.7µm while the diameters of non-primary dendrites were approximated by a regular tapering to reach a final diameter of 0.5µm. Firing frequency and AP analysis (amplitude, first and second derivative of APs) were computed online by handmade routines directly written in NEURON hoc language (Hines and Carnevale, 2001).

All computing files will be available at model DB database.

### Experimental design and statistical analysis

Statistical analysis (performed according to data distribution) included: linear regression, paired t-test, one-way ANOVA and Kruskal Wallis tests, with a p value <0.05 being considered statistically significant. For comparison between the four developmental stages, depending on the distribution of the data, we used a one-way parametric ANOVA followed by post hoc Tukey test for multiple comparisons or a non-parametric Kruskal Wallis with post-hoc Dunn test for multiple comparisons. Linear regressions were obtained with the Pearson test. Unless otherwise stated, statistical data are given as mean ± standard deviation and n indicates the number of analyzed neurons. Statistical tests were computed by using Sigmaplot 11.0 software (Systat Software Inc., San Jose, CA, USA) and Prism 6 (GraphPad Software, Boston, MA, USA).

### Figure preparation

Figures were prepared using Sigma Plot, Adobe Photoshop and Adobe Illustrator (Adobe Creative Cloud 2020, Adobe Systems Inc., San Jose, CA, U.S.A.), and ImageJ (Schindelin et al., 2012; Schneider et al., 2012; Rueden et al., 2017), with brightness and contrast adjustments performed consistently across the images to enhance clarity.

## Results

To investigate the relationship between changes in electrophysiological phenotype and morphology of SNc DA neurons during post-natal development, we performed patch-clamp recordings combined with post-hoc neuronal reconstructions based on neurobiotin fills of the recorded neurons at 4 different ages (post-natal day 3 or P3, n=14; P7, n=31; P14, n=13; P21, n=40, total n=98). These particular ages were chosen based on our previous description of the developmental trajectory of the electrophysiological phenotype of SNc DA neurons, which suggested that the two main transitions in electrical behavior occur respectively between P3 and P7 and between P7 and P14, P21 being considered a mature stage where SNc DA neurons produce the same electrophysiological phenotype as adult neurons (Dufour et al., 2014). Consistent with these choices, early studies of the morphological development of rat SNc DA neurons already suggested that basic morphological properties (soma length, dendrite diameter) reach maturity by P14 (Tepper et al., 1994; Park et al., 2000). Recordings were obtained from coronal midbrain slices in the presence of picrotoxin and kynurenate to block synaptic activity and isolate the intrinsic properties of SNc DA neurons. AnkyrinG immunostainings were performed in parallel of streptavidin staining to unambiguously identify the axon and measure axon initial segment (AIS) geometry. As can be seen on the representative pictures presented in Figure 1A, the general morphology of SNc DA neurons evolved strongly during the first three post-natal weeks. In particular, soma volume and the distance between the AIS and the soma (AIS-soma distance) strongly increased between P3 and P21 (Figure 1B).

**Figure 1.**
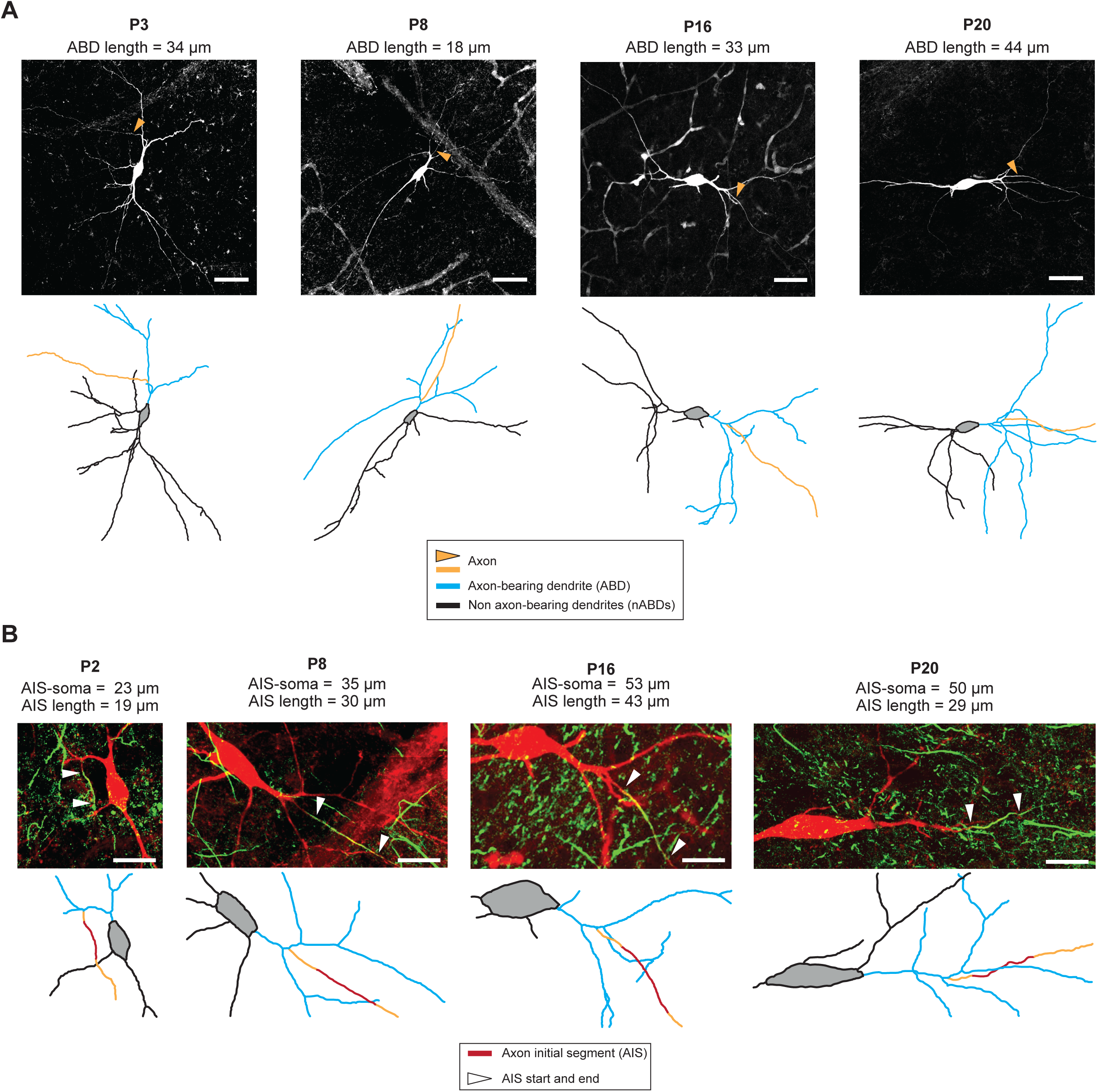
Establishment of neuronal morphology between P3 and P21. **A**, neurobiotin staining and skeleton representations of SNc DA neurons at P3, P8, P16 and P20. Regardless of age, most neurons exhibit an axon arising from an ABD, even though ABD length increases from P3 to P21. The number of ABD secondary dendrites also increases between P3 and P21. B, double neurobiotin-ankyrinG staining and skeleton representations of SNc DA neurons at P2, P8, P16 and P20 illustrating the maturation of AIS length and distance from the soma. AIS distance from the soma increases from P3 to P21. Each panel shows the skeleton with the original image at the same scale. Scale bars: **A**, 50µm; **B**, 25µm.

We then proceeded to a careful quantitative analysis of all morphological properties to determine which parameter changed significantly between these four developmental stages, and whether the timecourses of these changes were consistent (Figure 2). Based on our previous findings (Moubarak et al., 2022), we paid particular attention to the differential morphology of the axon-bearing and non axon-bearing dendrites (ABD and nABDs) and the geometry of the AIS. The parameters we measured are presented in Figure 2A: we defined dendritic complexity as the number of dendritic segments (a segment separates two branching points), and quantified it for the entire dendritic tree but also separately for the ABD and nABDs. We measured the total length of the entire dendritic tree, of the ABD and nABDs, the distance between the soma and the AIS, the length of the axonal segment preceding the AIS (axon start) and AIS length. We also calculated soma volume for each developmental stage. As every SNc DA neuron carries several nABDs, and in order to compare the properties of nABDs with the ABD, we calculated the average length and complexity of the nABDs for each neuron.

**Figure 2.**
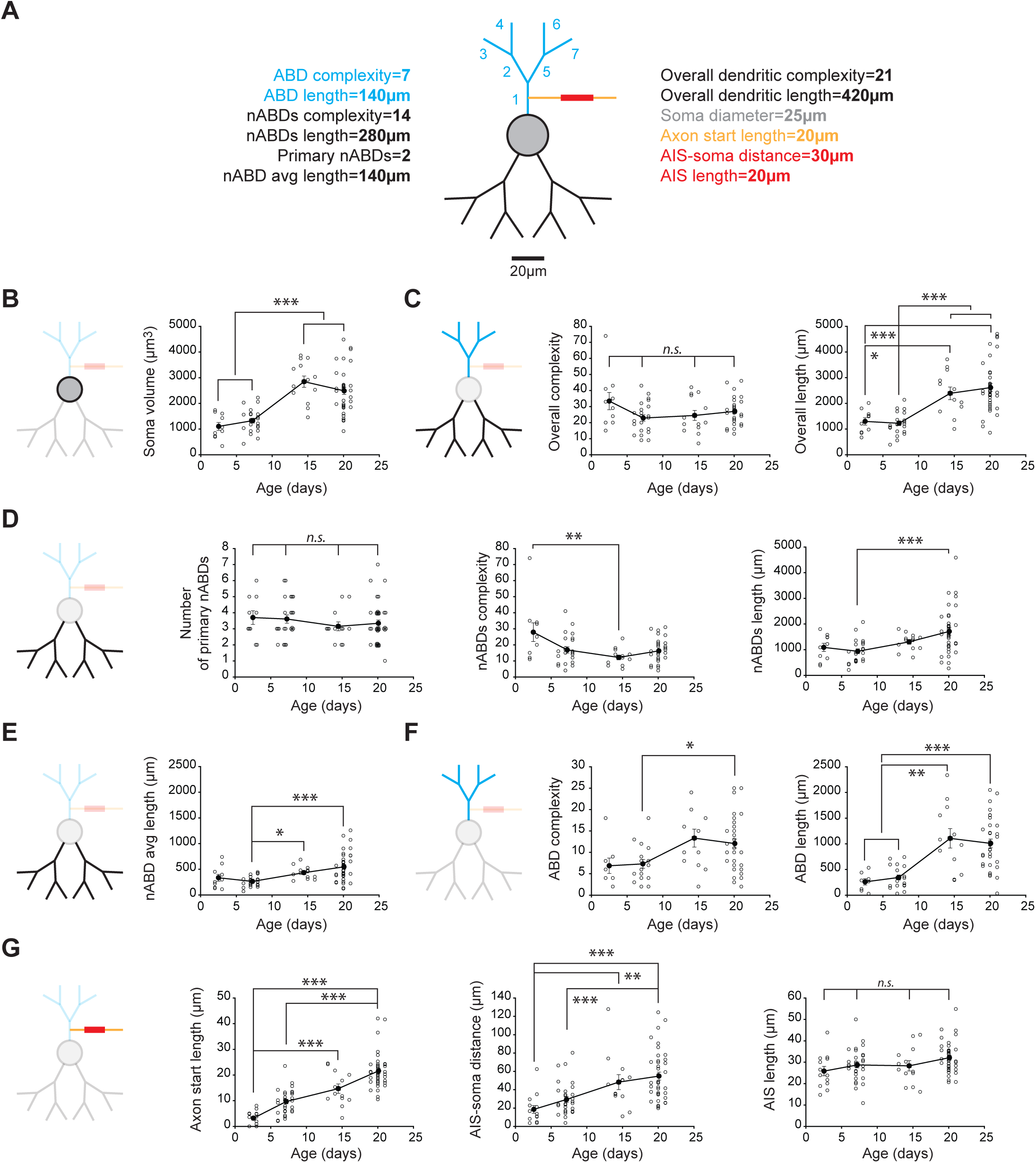
Postnatal developmental timecourse of morphological properties of SNc DA neurons. **A**, schematized fictive SNc DA neuron illustrating the measurements that have been performed to characterize morphology. The ABD is depicted in blue, while the nABDs appear in black. Soma is represented in grey, axon in orange, and AIS in red. Complexity was measured as the number of segments comprising a given dendrite, while primary dendrite length corresponded to the sum of length of these segments. In the following panels, a reduced version of the schematic is used to highlight the compartment analyzed. **B**, scatter plot showing the evolution of soma volume from P3 to P21. **C**, left, scatter plot showing the evolution of overall dendritic complexity from P3 to P21. Right, scatter plot showing the evolution of overall dendritic length from P3 to P21. **D**, left, scatter plot showing the evolution of the number of primary nABDs from P3 to P21. Middle, scatter plot showing the evolution of nABDs complexity from P3 to P21. Right, scatter plot showing the evolution of nABDs length from P3 to P21. **E**, scatter plot showing the evolution of nABD average length from P3 to P21. **F**, left, scatter plot showing the evolution of ABD complexity from P3 to P21. Right, scatter plot showing the evolution of ABD length from P3 to P21. **G**, left, scatter plot showing the evolution of axon start length from P3 to P21. Middle, scatter plot showing the evolution of the distance between the AIS and the soma from P3 to P21. Right, scatter plot showing the evolution of AIS length from P3 to P21. In each graph, open circles correspond to individual data points while closed dark circles represent the average for each developmental stage (P3, P7, P14, P21). Error bars correspond to SEM. Asterisks indicate statistical significance: non-significant n.s., p<0.05 *, p<0.01 **, p<0.001 ***.

We first looked at the evolution of the overall morphology. We found that soma volume was fairly stable between P3 and P7 (P3, 1103 ± 480 µm^3^, n=10 *vs* P7, 1328 ± 448 µm^3^, n=23) and between P14 and P21 (P14, 2845 ± 789 µm^3^, n=13 *vs* P21, 2489 ± 846 µm^3^, n=38), but abruptly increased between P7 and P14, such that soma volume at both P3 and P7 was significantly different from soma volume at both P14 and P21 (p<0.001) (Figure 2B, **Table 1**). The overall dendritic complexity was highly variable from cell to cell but the averaged complexity was fairly constant between P3 and P21 (no statistically significant difference between the four ages), with an average number of dendritic segments only showing a slight tendency to decrease after P3 (P3, 33.4 ± 17.02, n=10; P7, 22.87 ± 9.46, n=23; P14, 24.46 ± 11.34, n=13; P21, 27.03 ± 8.22, n=38) (Figure 2C, **Table 1**). Although we had previously described that dendritic length and complexity tend to be correlated with each other in P21 neurons for both the ABD and nABDs (Moubarak et al., 2022), the overall length of the dendritic tree showed strong changes during post-natal development (Figure 2C): similar to what has been observed for soma volume, no statistical differences were found between P3 and P7 (P3, 1297 ± 450 µm, n=10 vs P7, 1225 ± 430 µm, n=23) or between P14 and P21 (P14, 2392 ± 854 µm, n=12 vs P21, 2614 ± 957 µm, n=38), while P3 length was significantly shorter compared to P14 (p<0.05) and P21 (p<0.001) and P7 length was also significantly shorter than P14 and P21 lengths (p<0.001) (Figure 2C, **Table 1**).

**Table 1.** Summary of the statistics of the electrophysiological and morphological measurements performed on each developmental stage.

|  | P3 |  |  |  |  |  |  | P7 |  |  |  |  |  |  | P14 |  |  |  |  |  |  | P21 |  |  |  |  |  |  |
| --- | --- | --- | --- | --- | --- | --- | --- | --- | --- | --- | --- | --- | --- | --- | --- | --- | --- | --- | --- | --- | --- | --- | --- | --- | --- | --- | --- | --- |
|  | Mean | SD | Median | 25% | 75% | range | n | Mean | SD | Median | 25% | 75% | range | n | Mean | SD | Median | 25% | 75% | range | n | Mean | SD | Median | 25% | 75% | range | n |
| AP threshold (mV) | -44.57 | 3.60 | -45.65 | -47.13 | -42.80 | 12.22 | 14 | -44.59 | 3.92 | -44.94 | -47.19 | -41.43 | 14.61 | 34 | -43.97 | 5.24 | -42.68 | -47.10 | -40.09 | 19.17 | 13 | -44.68 | 3.53 | -44.97 | -47.69 | -41.87 | 18.20 | 54 |
| AP amplitude (mV) | 51.47 | 4.71 | 51.56 | 47.60 | 54.39 | 16.44 | 14 | 53.69 | 7.22 | 52.97 | 49.18 | 58.67 | 31.34 | 34 | 58.70 | 7.33 | 58.11 | 52.78 | 63.22 | 25.25 | 13 | 62.43 | 7.18 | 62.02 | 57.10 | 68.87 | 29.79 | 54 |
| AP half-width (ms) | 2.69 | 0.47 | 2.78 | 2.39 | 3.04 | 1.75 | 14 | 2.34 | 0.51 | 2.18 | 200.00 | 2.76 | 1.99 | 34 | 1.59 | 0.30 | 1.54 | 1.31 | 1.86 | 0.91 | 13 | 1.30 | 0.19 | 1.26 | 1.19 | 1.40 | 0.92 | 54 |
| IS dv2 peak (mV/ms2) | 122.47 | 42.52 | 114.14 | 94.91 | 159.95 | 136.64 | 14 | 123.06 | 33.54 | 123.63 | 99.29 | 136.12 | 170.07 | 34 | 311.96 | 109.61 | 313.38 | 215.51 | 361.85 | 373.65 | 13 | 178.28 | 54.72 | 165.10 | 140.14 | 216.62 | 241.06 | 51 |
| SD dv2 peak (mV/ms2) | 47.86 | 33.73 | 39.21 | 21.33 | 67.63 | 128.63 | 14 | 78.27 | 68.73 | 58.68 | 27.97 | 104.27 | 274.33 | 34 | 221.71 | 155.65 | 184.87 | 101.01 | 254.77 | 546.47 | 13 | 204.94 | 105.09 | 210.36 | 135.94 | 267.81 | 582.73 | 53 |
| SD/IS dv2 ratio | 0.44 | 0.32 | 0.36 | 0.16 | 0.67 | 1.04 | 14 | 0.62 | 0.47 | 0.51 | 0.28 | 0.90 | 1.72 | 34 | 0.73 | 0.48 | 0.52 | 0.42 | 0.92 | 1.73 | 13 | 1.26 | 0.83 | 1.09 | 0.71 | 1.54 | 4.66 | 50 |
| Overall dendritic segments | 33.40 | 17.02 | 31.00 | 20.00 | 41.00 | 59.00 | 10 | 22.87 | 9.46 | 23.00 | 15.25 | 29.50 | 34.00 | 23 | 24.46 | 11.34 | 20.00 | 16.50 | 37.25 | 35.00 | 13 | 27.03 | 8.22 | 27.00 | 22.00 | 33.00 | 33.00 | 38 |
| Segments on the ABD | 6.88 | 5.06 | 5.00 | 4.00 | 8.50 | 16.00 | 8 | 7.32 | 4.45 | 7.00 | 4.00 | 10.50 | 16.00 | 19 | 13.33 | 7.24 | 12.50 | 7.50 | 19.00 | 22.00 | 12 | 12.06 | 6.09 | 12.00 | 7.00 | 15.00 | 23.00 | 34 |
| Segments on nABDs | 27.90 | 18.49 | 22.50 | 15.00 | 34.00 | 63.00 | 10 | 16.83 | 9.35 | 15.00 | 8.25 | 23.75 | 37.00 | 23 | 12.15 | 5.38 | 12.00 | 7.00 | 14.75 | 19.00 | 13 | 16.24 | 6.76 | 16.00 | 12.00 | 20.00 | 27.00 | 38 |
| Number of primary nABDs | 3.70 | 1.34 | 3.50 | 3.00 | 5.00 | 4.00 | 10 | 3.61 | 1.23 | 3.00 | 3.00 | 4.75 | 4.00 | 23 | 3.15 | 0.99 | 3.00 | 2.75 | 3.25 | 3.00 | 13 | 3.34 | 1.28 | 3.00 | 2.00 | 4.00 | 6.00 | 38 |
| Avg. segment nbr per nABD | 8.26 | 5.66 | 5.80 | 4.00 | 12.50 | 15.17 | 10 | 4.56 | 1.71 | 4.50 | 4.00 | 5.25 | 6.40 | 23 | 4.13 | 2.29 | 3.00 | 2.46 | 5.75 | 6.67 | 13 | 5.37 | 3.15 | 5.00 | 3.00 | 6.33 | 17.00 | 38 |
| Total dendritic length (μm) | 1297 | 450 | 1349 | 876 | 1579 | 1342 | 10 | 1225 | 430 | 1179 | 945 | 1565 | 1748 | 23 | 2392 | 854 | 2344 | 1775 | 3098 | 2711 | 12 | 2614 | 957 | 2438 | 2021 | 3002 | 3853 | 38 |
| ABD length (μm) | 259 | 175 | 276 | 104 | 377 | 506 | 8 | 345 | 214 | 334 | 185 | 545 | 711 | 19 | 1106 | 656 | 1029 | 547 | 1634 | 2043 | 12 | 1008 | 495 | 982 | 604 | 1354 | 2014 | 34 |
| Total nABD length (μm) | 1090 | 505 | 1156 | 664 | 1489 | 1419 | 10 | 940 | 456 | 903 | 560 | 1258 | 1867 | 23 | 1306 | 327 | 1375 | 1060 | 1493 | 1110 | 13 | 1712 | 881 | 1602 | 1133 | 2021 | 4314 | 38 |
| Average length of nABDs (μm) | 335 | 213 | 275 | 145 | 451 | 629 | 10 | 263 | 99 | 259 | 189 | 307 | 376 | 23 | 433 | 123 | 425 | 339 | 492 | 416 | 13 | 547 | 274 | 521 | 376 | 689 | 1138 | 38 |
| Axon-soma distance (μm) | 15.38 | 17.33 | 12.40 | 3.40 | 17.79 | 62.51 | 14 | 19.73 | 18.47 | 16.11 | 8.24 | 23.51 | 74.58 | 31 | 33.59 | 27.64 | 25.01 | 17.81 | 47.92 | 103.35 | 13 | 33.25 | 27.54 | 28.39 | 9.69 | 51.90 | 105.91 | 40 |
| Axon start length (μm) | 3.20 | 2.73 | 3.68 | 0.00 | 4.95 | 8.06 | 14 | 9.68 | 4.87 | 10.00 | 5.86 | 13.03 | 21.44 | 31 | 14.72 | 6.54 | 12.69 | 10.65 | 20.50 | 21.25 | 13 | 21.61 | 6.70 | 20.88 | 17.54 | 23.90 | 31.48 | 40 |
| AIS-soma distance (μm) | 18.58 | 15.79 | 15.18 | 8.26 | 22.81 | 58.90 | 14 | 29.41 | 17.97 | 24.66 | 16.09 | 37.29 | 76.56 | 31 | 48.31 | 29.60 | 45.71 | 33.33 | 55.73 | 116.91 | 13 | 54.85 | 27.14 | 47.63 | 34.99 | 70.81 | 104.64 | 40 |
| AIS length (μm) | 25.85 | 7.98 | 23.91 | 19.77 | 31.63 | 29.14 | 13 | 28.74 | 7.60 | 28.49 | 24.34 | 33.06 | 39.13 | 30 | 28.31 | 8.58 | 26.55 | 23.16 | 34.01 | 26.68 | 13 | 32.14 | 7.14 | 30.67 | 27.58 | 35.55 | 34.22 | 40 |
| Soma volume (μm3) | 1103 | 480 | 996 | 727 | 1603 | 1399 | 10 | 1328 | 448 | 1324 | 1045 | 1573 | 1800 | 23 | 2845 | 789 | 2918 | 2301 | 3535 | 2408 | 13 | 2489 | 846 | 2510 | 1890 | 2724 | 3569 | 38 |

We then looked specifically at the properties of the nABDs. First, we found that the number of primary nABDs does not change significantly during post-natal development, with an average around 3.5 (Figure 2D, **Table 1**). However, the complexity of the nABDs appeared to slightly decrease during post-natal development (P3, 27.9 ± 18.49, n=10; P7, 16.83 ± 9.35, n=23; P14, 12.15 ± 5.38, n=13; P21, 16.24 ± 6.76, n=38), although the only significant difference was between P3 and P14 (p<0.01) (Figure 2D, **Table 1**). On the other hand, the total length of the nABDs appeared to slightly increase during post-natal development (P3, 1090 ± 505 µm, n=10; P7, 940 ± 456 µm, n=23; P14, 1306 ± 327 µm, n=13; P21, 1712 ± 881 µm, n=38), although the only significant difference was between P7 and P21 (p<0.001) (Figure 2D, **Table 1**). Because of the apparent opposite changes in complexity and length of the nABDs, and to compare with the single ABD, we calculated the average length of nABDs (Figure 2E, **Table 1**): average nABD length increased during post-natal development (P3, 335 ± 213 µm, n=10; P7, 263 ± 99 µm, n=23; P14, 433 ± 123 µm, n=13; P21, 547 ± 274 µm, n=38), with significant differences between P7 and both P14 (p<0.05) and P21 (p<0.001) (Figure 2E). However, the average number of segments per nABD did not change significantly with age: P3, 8.26 ± 5.66, n=10; P7, 4.56 ± 1.71, n=23; P14, 4.13 ± 2.29, n=13; P21, 5.37 ± 3.15, n=38 (**Table 1**).

The ABD displayed a developmental profile quite different from the nABDs (Figure 2F, **Table 1**). Although highly variable from cell to cell, ABD complexity appeared to increase with age (P3, 6.88 ± 5.06, n=8; P7, 7.32 ± 4.45, n=19; P14, 13.33 ± 7.24, n=12; P21, 12.06 ± 6.09, n=34), with P7 being significantly different from P21 (p<0.05) (Figure 2F, **Table 1**). ABD length displayed an even stronger increase with age: P3, 259 ± 175 µm, n=8; P7, 345 ± 214 µm, n=19; P14, 1106 ± 656 µm, n=12; P21, 1008 ± 495 µm, n=34 (Figure 2F, **Table 1**). The most dramatic change in length occurred between P7 and P14, such that P3 and P7 length were statistically different from P14 (p<0.01) and from P21 (p<0.001), while P3 length was not statistically different from P7 length nor was P14 length from P21 length.

Finally, we analyzed the post-natal development of the proximal axon, focusing on the AIS (Figure 2G, **Table 1**). First, the proportion of cells with an axon arising from the soma was fairly constant, with on average 13% of cells presenting this characteristic across the four developmental stages analyzed (2/10 at P3, 4/23 at P7, 1/13 at P14 and 4/38 at P21), suggesting that this general morphological feature of SNc DA neurons is defined very early in development. We then looked at the length of the axon start, which appeared to display a quasi-linear increase with age: P3, 3.20 ± 2.73 µm, n=14; P7, 9.68 ± 4.87 µm, n=31; P14, 14.72 ± 6.54 µm, n=13; P21, 21.61 ± 6.70 µm, n=40 (Figure 2G, **Table 1**). P3 axon start was significantly different from both P14 and P21 axon start (p<0.001) and P7 axon start was significantly different from P21 axon start (p<0.001). The distance between the AIS and the soma displayed a similar profile (P3, 18.58 ± 15.79 µm, n=14; P7, 29.41 ± 17.97 µm, n=31; P14, 48.31 ± 29.60 µm, n=13; P21, 54.85 ± 27.14 µm, n=40), with significant differences between P3 and both P14 (p<0.01) and P21 (p<0.001) and between P7 and P21 (p<0.001) (Figure 2G, **Table 1**). Unlike these two features, AIS length did not display any significant change across post-natal developmental stages: P3, 25.85 ± 7.98 µm, n=13; P7, 28.74 ± 7.60 µm, n=30; P14, 28.31 ± 8.58 µm, n=13; P21, 32.14 ± 7.14 µm, n=40 (Figure 2G, **Table 1**).

Overall, this analysis of soma, dendrites and proximal axon suggests that several morphological features of SNc DA neurons are set very early in development (overall dendritic complexity, number of nABDs, AIS length) while specific features display significant changes over the first three post-natal weeks, in particular between P7 and P14. To visualize the combined changes of all these morphological features, we built a scaled version of the average morphology of P3, P7, P14 and P21 SNc DA neurons (Figure 3A). In these scaled models, only soma diameter is represented on a non-linear scale (the represented soma diameter scales with the actual soma volume) in order to better visualize the significant changes between P7 and later stages. All other distances are scaled linearly with the changes observed in our morphological analysis (**Table 1**). Looking at these scaled models makes it immediately obvious that the most important changes in morphology occur between P7 and P14. Between these two stages, soma volume and ABD length abruptly increase to remain stable after P14. The differential change in morphology of the ABD and nABDs is even more evident when we look at the linearized representation of dendritic length (Figure 3A, right). In fact, this linearized version reveals that the ABD accounts for a small fraction of total dendritic length at P3 and P7 (19 and 27%, respectively), while it contributes almost as much as the nABDs at P14 and P21 (46 and 37%, respectively). This representation clearly shows that ABD and nABDs display strikingly different developmental timecourses. To pursue this investigation of the differences between ABD and nABDs, we then compared in each neuron the length and complexity of the ABD and the nABDs, using the average complexity and length for nABDs (Figure 3B). Comparing the number of segments per primary dendrite revealed that ABD and nABDs are equally complex at P3 (p=0.84), while the ABD is on average significantly more complex than the nABDs at P7 (p=0.02), P14 (p<0.001) and P21 (p<0.001). Similarly, ABD length is not significantly different from the averaged length of nABDs at P3 (p=0.55) and at P7 (p=0.15), while it is significantly longer at P14 (p=0.003) and at P21 (p<0.001). In summary, the ABD has properties very similar to nABDs until P7, but is significantly longer and more complex after P14.

**Figure 3.**
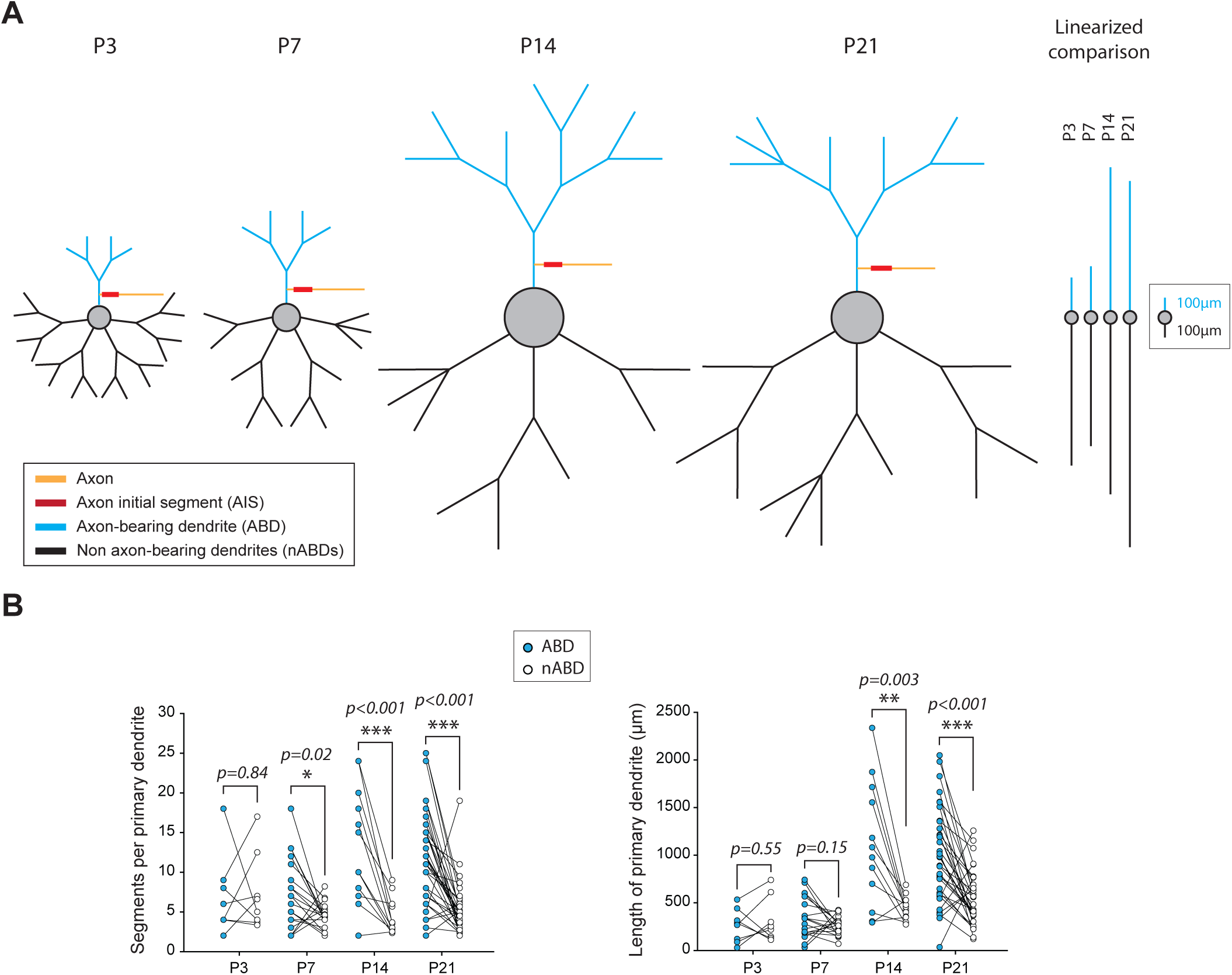
Postnatal development of dendritic asymmetry. **A**, left, scaled neuron models showing the average morphology of SNc DA neurons at P3, P7, P14 and P21. All distances (segment length, AIS length, AIS distance to the soma) have been scaled linearly. Soma diameter scales as a function of soma volume to better depict the abrupt change between P7 and P14. In order to better compare the kinetics of increase in length of ABD and nABDs, a linearized average of dendritic length is presented on the right (soma has not been scaled so that only dendritic length is visible). nABD length corresponds to the sum of all nABDs. The color coding is the same as the one presented in **Figure 2A**: ABD is in blue, nABDs in black, soma in grey, axon in orange and AIS in red. **B**, left, scatter plot illustrating the neuron-per-neuron comparison of ABD and averaged nABD complexity at P3, P7, P14 and P21. Right, scatter plot illustrating the neuron-per-neuron comparison of ABD and averaged nABD length at P3, P7, P14 and P21. Asterisks indicate statistical significance: p<0.05 *, p<0.01 **, p<0.001 ***.

To summarize these morphological analyses, the dendritic morphology of SNc DA neurons abruptly changes between P7 and P14, with a differential increase in length and complexity of the ABD compared to the nABDs. These results are consistent with the observations made in previous studies (Tepper et al., 1994; Park et al., 2000) showing that the overall morphology of SNc DA neurons was established by P14. Moreover, the dramatic change in morphology at this developmental stage (between P7 and P14, i.e. around P10) is highly reminiscent of the observations we made concerning the developmental trajectory of the electrophysiological phenotype of SNc DA neurons (Dufour et al., 2014).

Our previous work demonstrated that dendrites in mature SNc DA neurons contain a high density of sodium channels, and that this highly excitable dendritic compartment plays an essential role in the production of pacemaking but also in shaping the action potential recorded at the soma (Moubarak et al., 2019; Moubarak et al., 2022). Moreover, the latter study showed that the ABD and nABDs have opposite influences on AP duration, probably due to a slightly higher density of sodium channels in the ABD (Moubarak et al., 2022). Thus, we sought to determine whether developmental variations in AP shape in our reconstructed neurons were consistent with the observed morphological changes (Figure 4). In our previous studies, we showed that AP duration was determined by dendritic topology, and that using the second time-derivative of the voltage allowed a clear identification and measurement of the initial segment (IS) and somatodendritic (SD) components of the AP (Moubarak et al., 2019; Moubarak et al., 2022). Figure 4A presents the spontaneous activity and AP shape of neurons representative of each developmental stage, suggesting that AP duration decreases from P3 to P21, associated with differential changes in the contribution of the IS and SD components of the AP. Indeed, the statistical analysis showed that AP half-width decreases during development (P3, 2.69 ± 0.47 ms, n=14; P7, 2.34 ± 0.51 ms, n=34; P14, 1.59 ± 0.3 ms, n=13; P21, 1.3 ± 0.19 ms, n=54), with AP half-width at P3 and P7 being significantly longer than both P14 (p<0.01) and P21 (p<0.001) (Figure 4B, **Table 1**). Unlike AP half-width, both the IS and SD components of the AP did not follow a linear evolution (Figure 4B). Similar to what has been observed for several morphological properties, IS and SD components displayed an abrupt change between P7 and P14. The IS peak increased during development (P3, 122.47 ± 42.52 mV.ms^-2^, n=14; P7, 123.06 ± 33.54 mV.ms^-2^, n=34; P14, 311.96 ± 109.61 mV.ms^-2^, n=13; P21, 178.28 ± 54.72 mV.ms^-2^, n=51), with significant differences between P3 and both P14 (p<0.001) and P21 (p<0.01) and between P7 and both P14 (p<0.001) and P21 (p<0.001) (Figure 4B, **Table 1**). The SD peak displayed a similar increase (P3, 47.86 ± 33.73 mV.ms^-2^, n=14; P7, 78.27 ± 68.73 mV.ms^-2^, n=34; P14, 221.71 ± 155.65 mV.ms^-2^, n=13; P21, 204.94 ± 105.09 mV.ms^-2^, n=53), with significant differences between P3 and both P14 (p<0.01) and P21 (p<0.001) and between P7 and both P14 (p<0.01) and P21 (p<0.001) (Figure 4B, **Table 1**). Although both IS and SD peaks increased with a similar timecourse, the SD component appeared to display a larger increase, such that the overall SD/IS ratio displayed a significant increase during postnatal development (P3, 0.44 ± 0.32, n=14; P7, 0.62 ± 0.47, n=34; P14, 0.73 ± 0.48, n=13; P21, 1.26 ± 0.83, n=50), with a significant difference between both P3 and P7 and P21 (p<0.001) (Figure 4B, **Table 1**). In one of our earlier work (Moubarak et al., 2019), we showed that in mature neurons (P21), AP amplitude is mainly determined by the contribution of somatodendritic sodium channels, which is reflected by a strong correlation between SD peak and AP amplitude. In order to determine when in postnatal development this relationship is established, we looked at the correlation between the IS peak or SD peak and AP amplitude (Figure 4C). We found that, early in development (P3 and P7), AP amplitude is significantly correlated with IS peak (r=0.694, p=0.006, n=14 at P3; r=0.561, p=6.10^-4^, n=34 at P7) while it is not significantly correlated with IS peak at later stages (P14 and P21) (Figure 4C). On the other hand, AP amplitude correlates with SD peak at P7 (r=0.854, p=1.10^-10^, n=34), P14 (r=0.6662, p=0.014, n=13) and P21 (r=0.799, p=7.10^-13^, n=53) but not at P3 (Figure 4C). Thus, these results suggest that somatodendritic sodium channels play an increasingly important role in defining AP shape during post-natal development, with a clear dominance established at P14.

**Figure 4.**
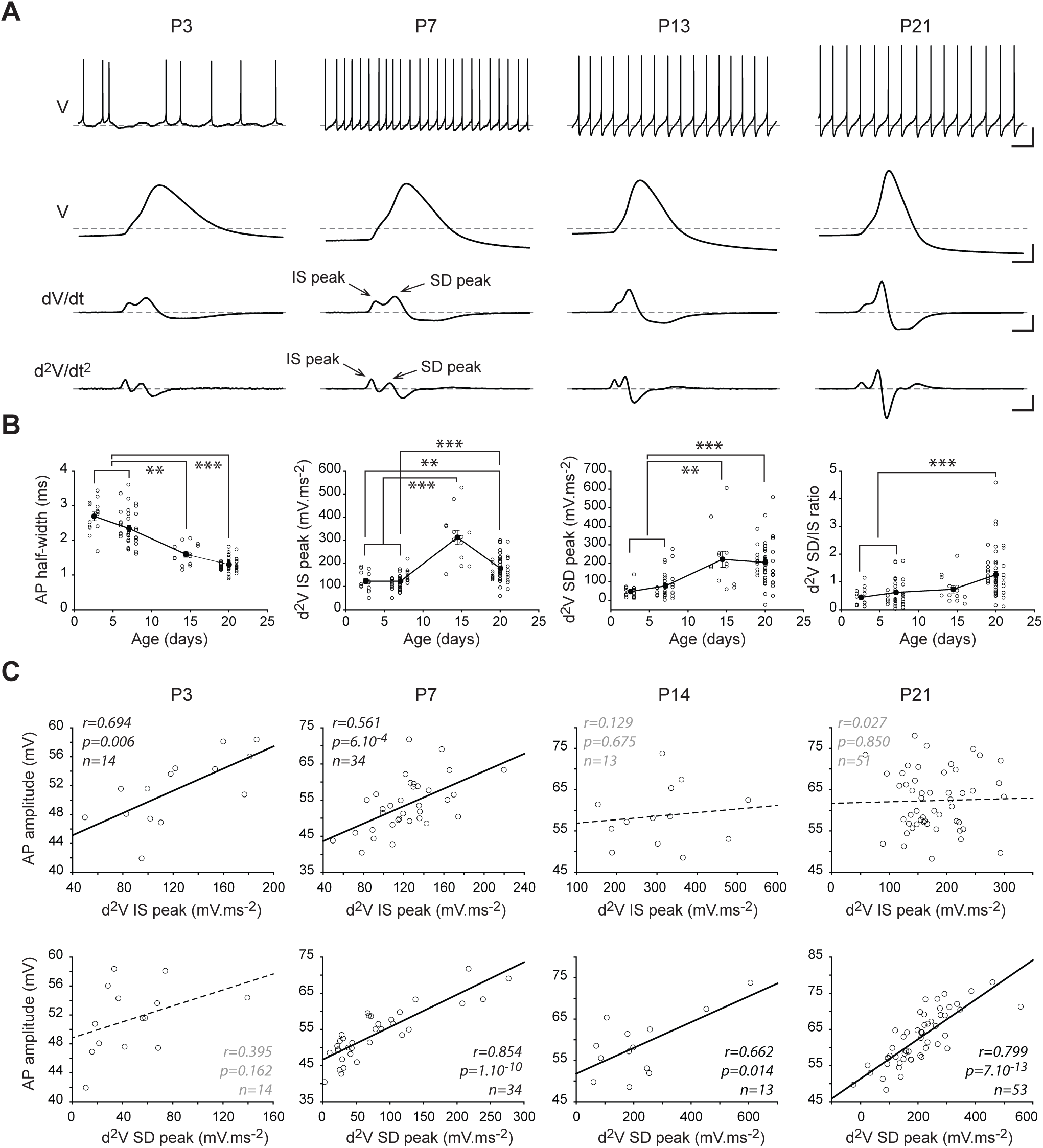
Postnatal development of action potential shape. **A**, representative traces showing the spontaneous activity pattern (top traces), the voltage signal during the AP on an expanded time scale (second from top), the first time-derivative of the voltage during the AP (second from bottom) and the second time-derivative of the voltage during the AP (bottom traces). The IS and SD peak on the first and second time-derivative of the voltage correspond to the initial segment and somatodendritic components of the AP. **B**, scatter plots representing the evolution of AP half-width (left), second time-derivative IS peak (second from left), second time-derivative SD peak (second from right) and SD/IS ratio (right) from P3 to P21. **C**, scatter plots showing the relationship between IS peak (top scatter plots) or SD peak (bottom scatter plots) and AP amplitude at each developmental stage (from P3 to P21, from left to right). Plain lines correspond to significant correlations while dotted lines indicate the absence of statistical significance. The r, p, and n values corresponding to the linear regression are presented on each scatter plot. Scale bars: **A**, top trace, vertical 20mV, horizontal 1s; second traces from top, vertical 20mV, horizontal 1ms; second trace from bottom, vertical 50mV.ms^-1^, horizontal 1ms; vertical 2000mV.ms^-2^, horizontal 1ms. Asterisks indicate statistical significance: p<0.01 **, p<0.001 ***.

So far, we showed that several morphological and electrophysiological parameters display a peculiar developmental timecourse, with substantial changes occurring between P7 and P14. In total, 14 morphological parameters and 6 electrophysiological parameters related to AP shape were analyzed. In order to obtain a more global visualization of the changes of these 20 parameters, we used a stacked representation of the statistical differences (Figure 5A) (Dufour et al., 2014). This representation clearly illustrates that no major change occurs between P3 and P7 or between P14 and P21, while the most significant changes occur between P7 and P14. This observation was confirmed by running agglomerative hierarchical clustering (AHC) of the data (n=68 neurons) using the 12 parameters showing the most significant differences (electrophysiological parameters E2 to E6; morphological parameters M1-M3, M7, M9, M12, M14) (Figure 5B). Using an automatic threshold for the dissimilarity index, AHC returned two classes corresponding to the early developmental stages (average age=6) and the late developmental stages (average age=17.8), supporting that P3 and P7 are very similar to each other and that P14 and P21 are also very similar to each other, both in terms of morphology and electrophysiological properties. Finally, we performed linear discriminant analysis (LDA), using 9 morphological and electrophysiological parameters to obtain a 2-dimensional representation of the morpho-electrical development of SNc DA neurons (Figure 5C). LDA provided results consistent with the statistical stacking and AHC analyses, showing that P3 and P7 cluster in one region of the F1/F2 space remote from the location of P14 and P21.

**Figure 5.**
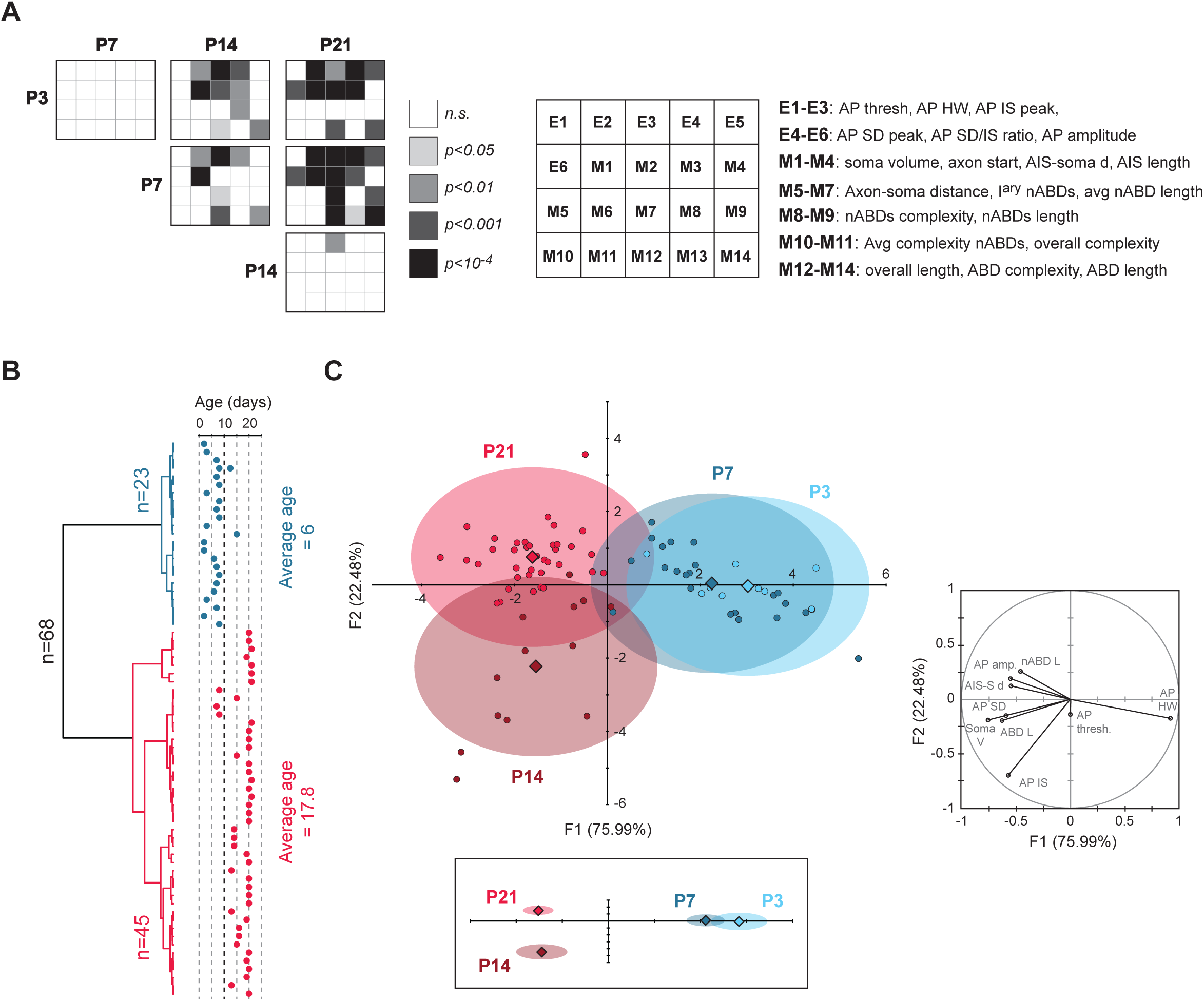
Multivariate analysis of the development of morphology and AP. **A**, statistical stacking table summarizing the statistical differences in 20 parameters (electrophysiological and morphological) between P3, P7, P14 and P21 stages. Each major square corresponds to the comparison between two developmental stages (column and row), while each subcell corresponds to one specific parameter (see legend on the right). The color of each subcell indicates the significance level of the difference found for a given parameter. None of the parameters are significantly different between P3 and P7 and only one parameter (IS peak) displays a statistical difference between P14 and P21. On the other hand, numerous statistical differences are found when comparing P3 or P7 to either P14 or P21. **B**, dendrogram representing the agglomerative hierarchical clustering of 68 neurons based on the 12 electrophysiological and morphological parameters showing the most significant differences across developmental stages: AP half-width, AP IS peak, AP SD peak, SD/IS ratio, AP amplitude, soma volume, axon start, AIS-soma distance, average nABD length, nABDs length, overall length, ABD length. The age corresponding to each neuron is represented on the right and shows that the two classes (blue and red) indeed correspond to early (P3-P7, average age P6) and late (P14-P21, average age 17.8) developmental classes. **C**, linear discriminant analysis (LDA) based on 9 electrophysiological and morphological parameters was performed on 82 neurons from P3, P7, P14 and P21. The left plot illustrates the distribution of the four age groups in the F1/F2 parameter space, showing that early stages (P3, P7, blue shades) strongly overlap in their distribution while they display negligible overlap with the late stages (P14, P21, red shades). Circles correspond to individual data points while the diamonds correspond to the centroid of each group. The lower plot (inset) shows the location of the four centroids (with the ellipses depicting the F1 and F2 SEM) in a scaled parameter space (based on F1 and F2 respective weights), emphasizing the proximity of P3 and P7 on one side and P14 and P21 on the other side. The polar plot on the right illustrates the relative contribution of the 9 electrophysiological and morphological parameters to F1 and F2.

To summarize, univariate and multivariate analyses of the morphological and electrophysiological properties (focused on AP shape) demonstrate that SNc DA neurons achieve their mature morphology and AP shape by P14, and that the most significant changes in morphology and AP shape occur between P7 and P14, such that SNc DA neurons from developmental stages earlier than P7 or later than P14 are difficult to distinguish from each other from both a morphological and electrophysiological viewpoints.

Since a number of experimental and theoretical studies including ours have suggested strong links between neuronal morphology and AP properties (Vetter et al., 2001; Kuba et al., 2006; Grubb and Burrone, 2010; Platkiewicz and Brette, 2010; Brette, 2013; Gulledge and Bravo, 2016; Kole and Brette, 2018; Moubarak et al., 2019; Moubarak et al., 2022), we then investigated the evolution of those interactions at each developmental stage. As the results for P3 and P7 (early stages) were very similar (data not shown), and the same was true for P14 and P21 (late stages), and the sample size was significantly larger for P7 (n=30) and P21 (n=39), only the results obtained for P7 and P21 are illustrated (Figure 6). The distance between the AP initiation site (i.e. the AIS in most neurons) and the soma is often described as having a major influence on the threshold and/or the kinetics of the AP around the threshold (also called sharpness, see (Platkiewicz and Brette, 2010; Brette, 2013; Kole and Brette, 2018)).

**Figure 6.**
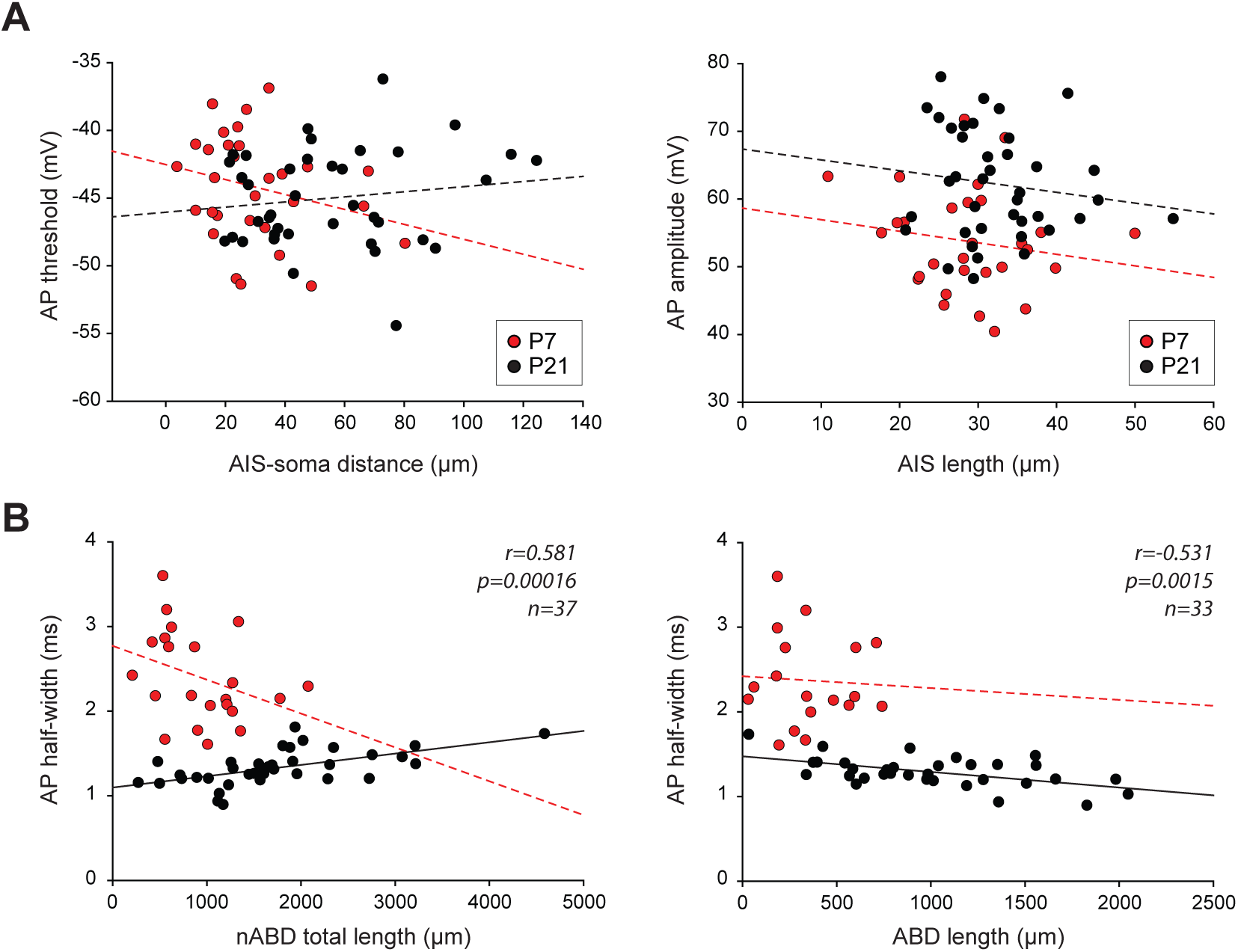
Relationships between morphology and AP shape in experimental data at P7 and P21. A, scatter plots showing the relationship between AIS-soma distance and AP threshold (left) and between AIS length and AP amplitude (right). B, scatter plots showing the relationship between nABD length (left) or ABD length (right) and AP half-width for P3, P7, P14 and P21 neurons. Dotted lines represent non-significant correlations while plain lines represent significant correlations. Please note that a significant correlation is present only for P21 neurons, with positive and negative correlations for nABDs and ABDs, respectively. The r, p, and n values corresponding to the statistically significant linear regressions are presented on each scatter plot.

Interestingly, we found that the AIS-soma distance was not significantly correlated with AP threshold at any developmental stage, including P7 (r=0.254, p=0.176, n=30) and P21 (r=0.144, p=0.380, n=39) (Figure 6A). Since our results showed that the IS component of the AP was significantly correlated with AP amplitude only at early stages while the SD component was most strongly correlated with AP amplitude at late stages (Figure 4C), we wondered whether AIS length (which should partly determine the total sodium conductance at the AIS) was correlated with AP amplitude (Figure 6A). Similar to AIS-soma distance, we found that AIS length was not correlated with AP amplitude at any developmental stage, including P7 (r=0.173, p=0.370, n=29) and P21 (r=0.141, p=0.393, n=39) (Figure 6A). Our most recent study demonstrated that, due to the presence of a high density of sodium channels in the somatodendritic compartment, AP half-width in mature SNc DA neurons (P21) is determined by dendritic topology, such that the ABD accelerates the AP while nABDs slow it down (Moubarak et al., 2022). These opposite influences were first spotted because ABD and nABD complexities (or lengths) displayed negative and positive correlations, respectively, with AP half-width. To determine whether the relationship between dendritic topology and AP half-width was present at early developmental stages (P3, P7, P14), we performed linear regressions using dendritic length (Figure 6B). A significant correlation between dendritic length and AP half-width was observed only at P21 (nABD, r=0.581, p=1.6.10^-4^, n=37; ABD, r=-0.531, p=0.0015, n=33) and absent from all the earlier developmental stages, including P7 (nABD, r=-0.347, p=0.114, n=22; ABD, r=-0.055, p=0.828, n=18) (Figure 6B).

The absence of relationship between morphology and AP shape at early developmental stages could be due to morphological development or to biophysical changes such as an increase in somatodendritic sodium channel density. To distinguish between these two hypotheses, we used realistic multicompartment Hodgkin-Huxley models of P7 and P21 SNc DA neurons based on the reconstructed morphology of the recorded neurons (Figure 7). Our previous work (Moubarak et al., 2022) showed that only models incorporating a higher density of sodium and calcium conductances in the ABD compared to the nABDs (g_Na_=120 *vs* 50pS/µm^2^, g_Ca_=2.2 *vs* 1pS/µm^2^) could replicate the relationship between dendritic topology and AP half-width observed in P21 neurons and illustrated in Figure 6B. Thus we built two models of P7 neurons, displaying either a homogeneous density of sodium and calcium conductances in the ABD and nABDs (Model #1) or a heterogeneous density of these conductances (Model #2) in the ABD and nABDs (Figure 7A). We then evaluated whether these models could replicate the observations presented in Figure 6. First we looked at the relationship between AIS and AP properties (Figure 7B). Surprisingly, we found that AIS-soma distance was weakly although significantly correlated with AP threshold for the Model #2 of P7 neurons (r=0.533, p=0.019, n=19) while no correlation was observed at P21 (r=0.193, p=0.259, n=36) (Figure 7B). No significant correlation was found for Model #1 at either age (data not shown). Then we looked at the relationship between AIS length and AP amplitude and found that these two parameters were not correlated in the P7 Model #2 (r=0.035, p=0.888, n=19) nor in the P21 Model #2 (r=-0.0251, p=0.884, n=36) (Figure 7B). Similar results were obtained for P7 and P21 Model #1. Thus, the model confirmed that AIS morphology does not seem to affect AP shape in early or late developmental stages, an observation consistent with our previous (Moubarak et al., 2019; Moubarak et al., 2022) and current experimental results (Figure 6).

**Figure 7.**
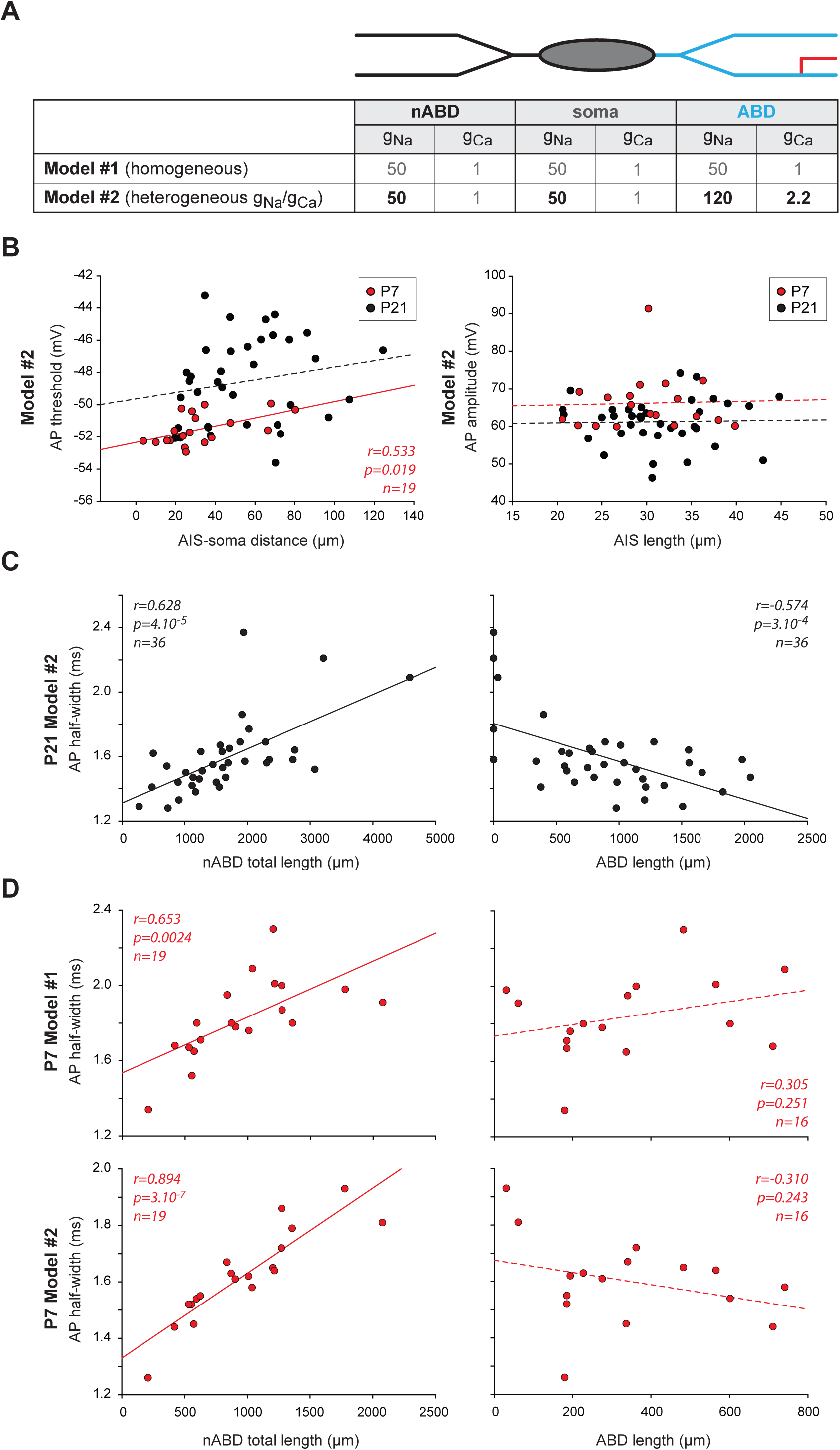
Relationships between morphology and AP shape in P7 and P21 multicompartment models. **A**, table presenting the distribution of the sodium (g_Na_) and calcium (g_Ca_) conductance densities (in pS/µm^2^) in the homogeneous (Model #1) and heterogeneous (Model #2) versions of the multi-compartment Hodgkin-Huxley models. **B**, left, scatter plot showing the relationship between AIS-soma distance and AP threshold in the heterogeneous version of the model of P7 and P21 neurons. Right, scatter showing the relationship between AIS length and AP amplitude in the heterogeneous version of the model of P7 and P21 neurons. Dotted lines represent non significant correlations while plain lines represent significant correlations. The r, p, and n values corresponding to the statistically significant linear regressions are presented on each scatter plot. **C**, scatter plots showing that the Model #2 of P21 neurons reproduces the positive and negative correlations observed between nABD or ABD length, respectively, and AP half-width. The r, p, and n values corresponding to the linear regressions are presented on each scatter plot. **D**, top, scatter plots showing the relationship between nABD length (left) or ABD length (right) and AP half-width in the Model #1 of P7 neurons. The r, p and n values are provided, showing that only nABD length is significantly correlated with AP half-width. Bottom, scatter plots showing the relationship between nABD length (left) or ABD length (right) and AP half-width in the Model #2 of P7 neurons. The r, p and n values are provided, showing that only nABD length is significantly correlated with AP half-width.

Then, we focused on the relationship between dendritic topology and AP half-width. Figure 7C presents the results obtained in our previous work (Moubarak et al., 2022) showing that the heterogeneous model of P21 neurons replicates the opposite relationships between the ABD and nABD lengths and AP half-width. To determine whether the absence of correlations between dendritic length and AP half-width observed at early developmental stages (Figure 6B) could be explained by changes in sodium channel density, we tested these correlations in both the homogeneous and heterogeneous models of P7 neurons (Figure 7D). Interestingly, while both the homogeneous and heterogeneous models of P7 neurons displayed a positive correlation between nABD length and AP half-width similar to the one observed in P21 neuron models (Figure 7C), the negative correlation between ABD length and AP half-width observed in heterogeneous models of P21 neurons was not observed in P7 models, neither for the homogeneous nor for the heterogeneous version of the model (Figure 7D). These results suggest that the strong relationship between dendritic topology and AP duration observed in mature neurons (Figure 6B, (Moubarak et al., 2022)) is due to a synergistic effect of dendritic morphology (ABD more complex than nABDs in P21 neurons) and increased sodium channel density.

To explore further this hypothesis, we used a slightly different screening approach with the computational models. In our previous work, we showed that the heterogeneous model of P21 neurons not only reproduced the relationship between dendritic topology and AP half-width but also predicted the half-width of the AP recorded in the real neurons (Moubarak et al., 2022). Since all model neurons had the same biophysical properties (g_Na_=120pS/µm^2^ and g_Ca_=2.2pS/µm^2^ in the ABD, g_Na_=50pS/µm^2^ and g_Ca_=1pS/µm^2^ in the nABDs) but distinct morphologies (neuronal reconstruction), this suggested that cell-to-cell variations in dendritic morphology were the main factor defining the cell-to-cell variations in AP half-width in mature neurons. Our current results suggest that P7 neurons might follow different rules, thus we sought to define the respective impact of morphology and sodium conductance density on AP half-width in P7 and P21 neurons. To do so, we tested a range of g_Na_ values for each model and selected the g_Na_ that yielded an AP half-width value most similar to the biological AP half-width. Since P21 heterogeneous models already predicted biological AP half-width (r=0.718, p=1.10^-6^, n=35, see (Moubarak et al., 2022)), we started from this model and only explored variations of g_Na_ in the ABD that would improve the correlation (g_Na_ was varied from 50 to 150pS/µm^2^ in steps of 10pS in the ABD, g_Ca_ being kept constant at 2.2pS/µm^2^). For P7 models though, we observed that heterogeneous models were unable to predict biological AP half-width (data not shown) and thus tested g_Na_ variations in the homogeneous model. In this latter case, g_Na_ was varied simultaneously in the ABD and nABDs between 25pS/µm^2^ and 165pS/µm^2^ in steps of 10pS. For both models, we then interpolated or extrapolated the values of g_Na_ that yielded the AP half-width most similar to the biological value. As can be seen in Figure 8A, the cell-by-cell tuning of g_Na_ allowed us to predict AP half-width in both the P7 and P21 models, although the correlation was stronger in P21 models. Interestingly, the optimal g_Na_ was quite different between P7 and P21 models (Figure 8B): the average optimal g_Na_ was 29.42 ± 23.04pS/µm2 in P7 homogeneous models (n=19) while it was 118.57 ± 37.24pS/µm2 in P21 heterogeneous models (n=35), with a significant statistical difference (p<0.001, Mann-Whitney test). These results suggest that g_Na_ might be much lower in P7 neurons compared to P21 neurons, reinforcing our hypothesis of a steep increase in g_Na_ during post-natal development. In addition, since every discrete value of g_Na_ was tested on each neuron, we could estimate the sensitivity of each neuron to changes in g_Na_ density by measuring the coefficient of variation of AP half-width for a fixed range of g_Na_. In order to compare the sensitivities of P7 and P21 models, we used the whole g_Na_ range for P21 models (50 to 150pS) and restricted the range of g_Na_ values to 55-165pS for P7 models, corresponding to a 3-fold range for both stages. Interestingly, P7 model neurons appeared more sensitive to g_Na_ variations than P21 models, as the CV was significantly larger for the P7 compared to P21 models (CV = 11.67 ± 3.27 for P7 *vs* 3.45 ± 2.57 for P21, p<0.001, Mann-Whitney test, Figure 8C). This result reinforces the idea that dendritic topology is the main factor defining the cell-to-cell variations in AP half-width in P21 neurons, while cell-to-cell variations of g_Na_ might predominate in P7 neurons. Altogether, the computational models confirm that the key influence of the dendritic arborization on AP shape observed in mature neurons might arise from a synergistic increase in dendritic length (and complexity) and sodium conductance density in the somatodendritic compartment.

**Figure 8.**
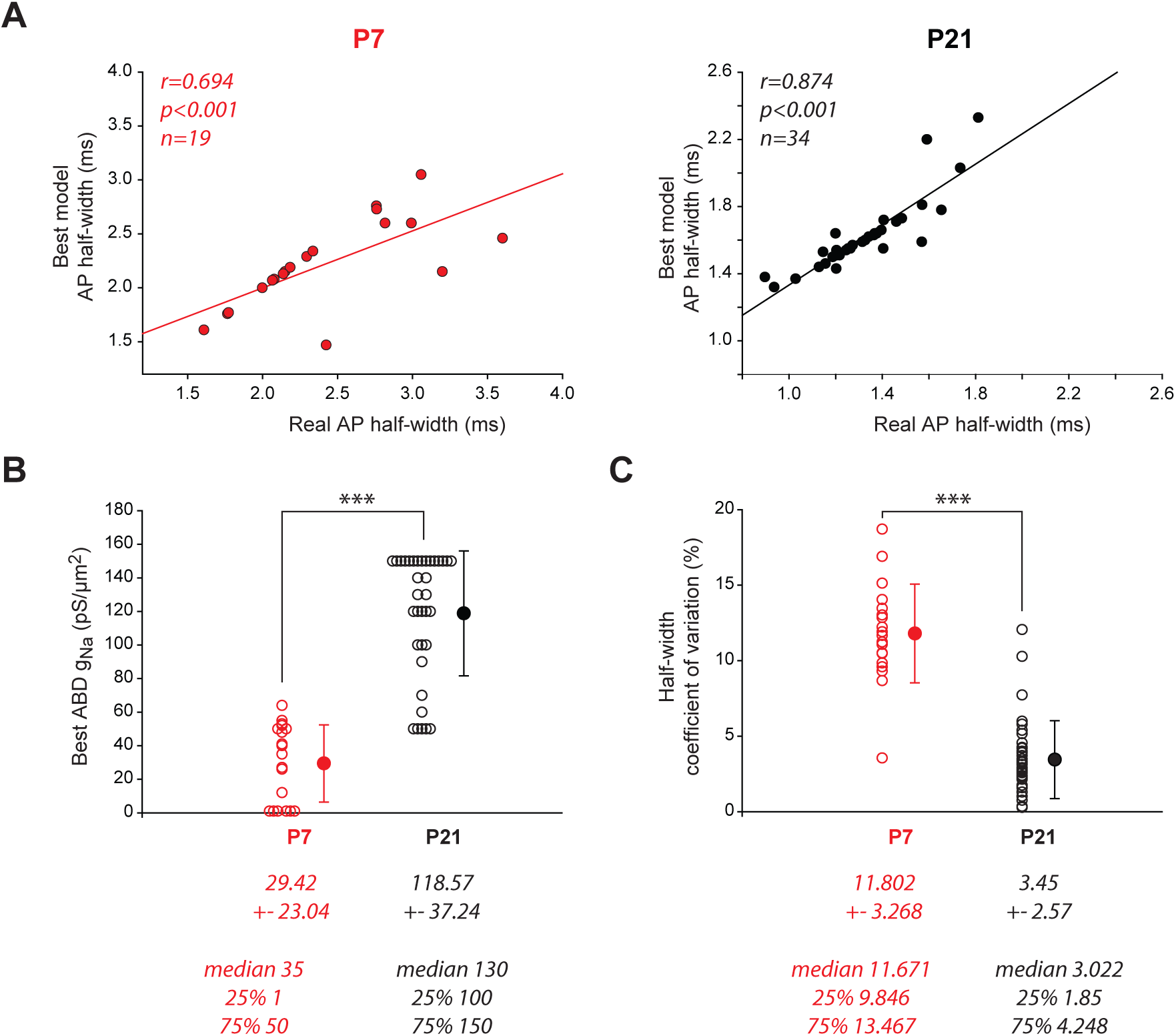
Determining the optimal sodium conductance in P7 and P21 multicompartment models. **A**, scatter plots representing the correlation between the values of the AP half-width recorded in the biological neurons and the best predicted values of AP half-width obtained in the corresponding realistic models for P7 (left) and P21 neurons (right). The r, p, and n values corresponding to the linear regressions are presented on each scatter plot. **B**, scatter plot representing the distribution of values of g_Na_ best predicting real AP half-width (same models as in panel **A**) in the P7 (red circles) and P21 models (black circles). Open circles correspond to individual data points while closed dark circles represent the average for each developmental stage. Error bars correspond to SD. **C**, scatter plot representing the distribution of the coefficients of variation of AP half-width associated with variations in g_Na_ in P7 (red circles) and P21 models (black circles). Open circles correspond to individual data points while closed dark circles represent the average for each developmental stage. Error bars correspond to SD. Asterisks: *** p<0.001.

## Discussion

In the present study, we demonstrated that the mature dendritic morphology of SNc DA neurons is acquired during the first three postnatal weeks, with the most significant changes occurring specifically between P7 and P14. In particular, the complexity and length of the ABD and nABDs differentially increase at this stage, such that the ABD becomes significantly longer and more complex than the other dendrites at P14. Soma volume also shows an abrupt increase between P7 and P14. In constrast, some morphological features (AIS distance from the soma for instance) display a quasi-linear increase over the first three postnatal weeks, and others do not change significantly (AIS length, number of primary dendrites), suggesting that dendritic morphology and soma volume are under the control of distinct regulatory processes. In parallel, and consistent with the demonstrated role of dendrites in shaping the AP (Moubarak et al., 2019; Moubarak et al., 2022), the IS and SD components of the AP also undergo abrupt changes in amplitude between P7 and P14 to remain stable thereafter. Consistent with our previous observations (Moubarak et al., 2019; Moubarak et al., 2022), we show that AIS morphology has little impact on AP shape at any postnatal developmental stage. In addition, we show that the influence of dendritic topology on AP shape is only present at mature stages (P21), and realistic computational modeling suggests that this relationship relies on the parallel increase in length and complexity of the ABD and the increase in somatodendritic sodium conductance occurring between P7 and P14. To summarize, the current study suggests that crucial biophysical and morphological changes occur between P7 and P14 to provide SNc DA neurons with their mature morphology and action potential shape by the end of the second postnatal week.

The reported change in soma volume is consistent with three previous studies (Lauder and Bloom, 1975; Tepper et al., 1994; Park et al., 2000) also reporting a substantial increase in soma size until P14 and a stability of this parameter at later developmental stages. Lauder and Bloom for instance reported that nucleus size increases until P15 and remains stable in later stages and adulthood (Lauder and Bloom, 1975). In addition, the study of Park et al. also showed that the length of DA terminals surrounding cell bodies in the lateral striatum increases until P17 but then stays fairly constant in the adult rat (at least until P75) (Park et al., 2000). Similar results were already obtained by older studies using light and electron microscopy (Hattori and McGeer, 1973; Voorn et al., 1988) that demonstrated that synaptic terminal density in the striatum increases rapidly during postnatal development, with the most significant increase occurring between P13 and P17, and marginal increases in density after these developmental stages (300g adult rats were also analyzed). Another observation that has been made by the group of Burke (Burke, 2003; Ries et al., 2009) is that, although most of DA neuron cell death occurs perinatally (between E18 and P8), a second peak of cell death is observed at the end of the second postnatal week (P14). Therefore, all these observations seem to indicate that major morphological maturation processes take place during the first two postnatal weeks in SNc DA neurons, reinforcing our conclusion that most of the changes in somatodendritic morphology occur before the end of the second postnatal week.

From a functional point of view, our results are reminiscent of the data that were obtained concerning the developmental trajectory of the electrophysiological phenotype of SNc DA neurons (Dufour et al., 2014). In this former study where we performed a more exhaustive screening of postnatal developmental stages (P2 to P29 in ∼2-day increments), we demonstrated that the firing pattern of rat SNc DA neurons is acquired in two sequences, with major electrophysiological changes occurring specifically between P3 and P5 and then between P9 and P11 before stabilizing after P14. Moreover, we demonstrated that the second transition (P9-P11) seems to be supported by an increase in sodium conductance and calcium-activated potassium conductances (SK channels). We also demonstrated that, after P14, electrophysiological phenotype is stable (at least until P29, the older age tested in those experiments). While these results suggested that an increase in sodium conductance between P9 and P11 was partly responsible for the maturation of the electrophysiological phenotype, we did not provide insights into the mechanisms responsible for this increase. On one side, the present results suggest that the rapid increase in length and complexity of the ABD is partly responsible for this increase in sodium conductance. Specifically, we found that ABD length is scaled up by a factor of at least 2.5-3 between P7 and P14, going from less than 350µm on average to more than 1000µm. Even without considering the probable associated increase in dendritic diameter or potential changes in the density of sodium channels expressed at the somatodendritic membrane, this means that the total ABD sodium conductance is likely to increase by a factor of at least 2.5-3 between P7 and P14. Thus, the increase in ABD length and complexity could explain a substantial part of the increase in sodium conductance that seems to underlie the acquisition of regular pacemaking (Dufour et al., 2014). On the other hand, the modeling results presented in Figure 8 show the somatodendritic sodium conductance densities required to predict AP half-width in P7 neurons and P21 neurons are drastically different (∼30pS/µm^2^ in P7 neurons compared to ∼130pS/µm^2^ in P21). While these modeling results are not definite proof, they suggest that sodium channel density may strongly increase between P7 and P14 in order to explain the change in AP shape. Overall these results lead us to conjecture that the changes in electrophysiological phenotype between P7 and P14 are driven by a synergistic increase in ABD length and in somatodendritic sodium channel density.

This hypothesis would need to be confirmed, using assays that directly quantify the expression level of sodium channels in specific compartments, e.g. immunohistochemistry. Unfortunately, immunostainings for sodium channels do not efficiently reveal somatodendritic expression, which is considerably weaker than at the AIS. A recent study using immunohistochemical stainings in mouse SNc DA neurons (Yang et al., 2019) could ascertain the presence of Nav1.2 at the AIS, but did not reveal somatodendritic expression of this sodium subunit. In the same study, immunohistochemistry failed to reveal the presence of Nav1.1 and Nav1.6 in SNc DA neurons, neither in the axon nor in the dendrites. Although SNc DA neurons exhibit a higher density of sodium current than most cell types in the somatodendritic compartment (Moubarak et al., 2019), our realistic computational model suggests that AIS sodium conductance density must be at least 10-15 times higher than somatodendritic density to faithfully reproduce the AP behavior (initiation at the AIS, amplitudes of IS and SD components, back-propagation) observed in real neurons (Moubarak et al., 2019; Moubarak et al., 2022). This difference in density likely explains why sodium channels are not detected in the dendrites using immunohistochemistry. Thus, our hypothesis will need further experiments to confirm that the changes in AP shape observed between P7 and P14 are indeed a consequence of the sole changes in dendritic morphology we described here or whether modifications in somatodendritic sodium channel density and/or subtype are involved. Nav1.2 is the main subunit expressed in mature SNc DA neurons (Yang et al., 2019) but nothing is known about the subunits expressed at early developmental stages (P3, P7). Other subunits are expressed at the mRNA level in SNc DA neurons (Nav1.1, Nav1.3, Nav1.6) (Ding et al., 2011; Tapia et al., 2018) and could be potentially expressed during early postnatal development. A recent study showed that Nav1.3 is reexpressed in the SNc 8 weeks after a substantial lesion induced by a 6-OHDA injection (Wang et al., 2018). This reexpression suggests that Nav1.3 might be expressed at early developmental stages, although this hypothesis needs to be tested. Alternatively, but not exclusively, the changes in morphology and electrophysiological phenotype between P7 and P14 might be related to an increase in synaptic activity impinging on SNc DA neurons. However, it seems that synaptic contacts in the SNc appear around birth and their number increases gradually until P60, with no indication that a particularly strong increase in synaptic inputs occurs around P10-P14.

Independent of the mechanism underlying these developmental changes in AP properties, our experimental and modeling results also suggest that the influence of morphology on AP shape is restricted to mature SNc DA neurons. First, in spite of the more compact morphology and the associated relative proximity of the AIS at early developmental stages (P3, P7), we did not observe any relationship between AIS morphology and AP shape at any age. This might seem surprising as many studies (Kuba et al., 2006; Grubb and Burrone, 2010; Kuba, 2012; Gulledge and Bravo, 2016; Kole and Brette, 2018; Meza et al., 2018) have shown that AIS morphology can have a profound impact on excitability in various neuronal types. However, our current results are consistent with our previous work (Moubarak et al., 2019; Moubarak et al., 2022), which demonstrated that the strong somatodendritic excitability arising from a high density of sodium channels considerably weakens the influence of the AIS on somatically recorded activity in SNc DA neurons. While the analysis of the AP waveform showed a strong correlation between the IS component and AP amplitude at P3 and P7, this link was not corroborated by a significant correlation between AIS morphology and AP threshold, amplitude or half-width. Computational modeling applied to P7 and P21 neurons gave consistent results, although AP threshold displayed a significant correlation with AIS-soma distance in the heterogeneous model of P7 neurons. Apart from this correlation, none of the AP properties were correlated with AIS morphology in P7 or P21 neurons. Most interestingly, the link between dendritic topology and AP half-width observed in mature neurons and replicated in P21 heterogeneous models (Figure 7C) (Moubarak et al., 2022) was not observed in younger developmental stages and could not be reproduced in the P7 models. More precisely, only the positive correlation between nABD and AP half-width was reproduced by the model, consistent with our previous results showing that this positive correlation is found in any version of the P21 neurons (Moubarak et al., 2022). The negative correlation between ABD length and AP half-width was not observed in both the homogeneous and heterogeneous versions of the P7 model. Since we applied the same distribution of sodium conductances in the P7 and P21 models, we assume that the increase in length of the ABD (*3.5 in P21 compared to P7 neurons) is responsible for the appearance of the influence of the ABD on AP half-width. In other terms, these results suggest that both an asymetric distribution of sodium conductance (slightly higher in the ABD, (Moubarak et al., 2022)) and a long and complex ABD are necessary to confer to the ABD its strong influence on somatic AP shape.

In conclusion, the results obtained in this study refine the observations made previously on the development of SNc DA neuron morphology and electrophysiological phenotype, and suggest that major (but so far unknown) regulatory events controlling dendritic growth, dendritic excitability, but also maybe axon growth in the striatum, occur around P10 and are responsible for the final maturation of SNc DA neurons. In addition, our results reinforce the idea that, although the AP is initiated at the AIS, dendrites, and in particular the ABD, play a central role in establishing and sustaining of the mature electrophysiological phenotype of SNc DA neurons. Future work will be necessary to determine the nature of the signals involved in the crucial morphological and electrophysiological developmental step described in this study.

## Author contributions

F.T. and J.M.G. conceptualized research. E.M., F.T. and J.M.G. designed the methodology. E.M., F.W. and F.T. performed research. E.M., F.T. and J.M.G. analyzed data. E.M., F.T. and J.M.G. wrote the manuscript original draft. Further writing, reviewing and editing was done by E.M., F.W. and J.M.G.

## Acknowledgements

this work was supported by the French Ministry of Research (doctoral fellowship to E.M.) and the European Research Council (Consolidator grant 616827 *CanaloHmics* to J.M.G.).

